# Characterization of Isoforms of the Ovine Granulocyte Colony Stimulating Factor

**DOI:** 10.1101/849166

**Authors:** Runting Li, Longxin Chen, Yuqin Wang, Limeng Zhang, Ting Liu, Xiaoning Nie, Haiying He, Yong Wang, Kang Wang, Ruochen Yang, Chunhui Duan, Yueqin Liu, Runlin Zhang Ma, Yingjie Zhang

## Abstract

The granulocyte colony-stimulating factor (GCSF) regulates the maturation, proliferation, and differentiation of precursor cells of neutrophilic granulocytes, and has been widely studied in several species. To investigate the function of variants of sheep GCSF (sGCSF), this study compared difference in their mRNA expression levels. Both the activity and mRNA expression level of GCSFv2 were higher than those of GCSFv1. Their sequences were aligned, which showed that they had the highest homology with bovine GCSF. Then, predicted ovine GCSF isoforms and their constant C-terminals were cloned and expressed, which were stably expressed in mammalian cells. After purification, all GCSF functions were different both *in vitro* and *in vivo*, and the GCSF C-terminal was best. These results indicated that the ability to stimulate both the proliferation and differentiation of progenitor cells and to activate the maturation of neutrophils could be used for research of efficacious non-antibiotic protein drugs. Furthermore, GCSF can be used as candidate target of genetic breeding to specifically improve sheep immunity.

## Introduction

The granulocyte colony-stimulating factor (GCSF) is the principal growth factor that regulates the maturation, proliferation, and differentiation of precursor cells of neutrophilic granulocytes (Nickerson 1991). The GCSF coding gene is localized on chromosome 11. The GCSF is a member of the long-chain subtype of the class 1 cytokine superfamily, which includes growth hormone, erythropoietin, interleukin 6, and oncostatin M. The crystal structure of the GCSF is complexed to the BN-BC domains, which is the principal ligand-binding region of the GCSF receptor (GCSFR), and forms a complex at a 2:2 ratio with the ligand. This complex has a non-crystallographic pseudo-twofold axis, which primarily runs through the interdomain region and secondarily the BC domain (Aritomi *et al*. 1999). The GCSF plays a role in regulating the phosphatidylinositol-3-kinase/serine/threonine kinase (PI3K/AKT) pathway, thus affecting cell proliferation and apoptosis, inducing the expression of the vascular endothelial growth factor (VEGF) (Liang *et al*. 2018).

Most antibiotics are used for livestock, which is continuously increasing due to the increased global demand for meat. A growing body of evidence has linked this practice with the increase of antimicrobial-resistant infections, not just in animals but also in humans (Van Boeckel *et al*. 2019). During bacterial infections, the GCSF is produced and accelerates neutrophil production from their progenitors (Takehara *et al*. 2019), which may be utilized to avoid the use of antibiotics. Administration of GCSF to pigs has been shown to result in a longer mean survival time after exposure to *Streptococcus suis* (Brockmeier *et al*. 2019). The outcomes of naturally canine papillomavirus infected pups treated with the recombinant canine granulocyte-colony stimulating factor (rcGCSF), in combination with routine therapy, were compared with similarly-managed infected pups that had not been treated with rcGCSF (Armenise *et al*. 2019).

The GCSF has been widely studied in several species (Armenise *et al*. 2019; Brockmeier *et al*. 2019; Du *et al*. 2019; Katakura *et al*. 2019; Li *et al*. 2019; Sameni *et al*. 2019). There are four transcript variants of the GCSF in the human genome, two transcript variants in the mice genome, and one transcript in the bovine genome. The recombinant human GCSF transcript variant B has been commercially available since 1992 (Fraser *et al*. 1994); however, several unanswered questions need to be addressed by clinical trials (Eapen *et al*. 2019; Nguyen 1994). The GCSF has been widely used for both the prophylaxis and treatment of neutropenia in cancer patients and also for the mobilization of peripheral blood stem cells (PBSC) (Wu *et al*. 2019). GCSFs decrease the incidence of febrile neutropenia (FN) in patients who receive myelosuppressive chemotherapy upon FN incidence (Stephens *et al*. 2019). Only two predicted gene sequences of sheep granulocyte-colony stimulating factor (sGCSF) can be found in GenBank.

So far, it remains unknown whether sGCSF could be used to improve sheep immunity as has been achieved for other species. To clarify the function of sGCSF variants, their expression differences were compared at mRNA level to ensure their existence. Then, their sequences were aligned, the sequences of sGCSF variants 1 and 2, and their constant C-terminal were cloned to construct expression plasmids for expression and purification. The purified proteins were used for an *in vitro* bioassay and an *in vivo* test to study their functions.

## Materials and methods

### Vectors, cells, and animals

The pRTL1 plasmid for the generation of a GCSF eukaryotic expression vector was provided by Zhengzhou Normal University, China. M-NFS60 cells were kindly provided by Dr. Feng Wang (Institute of Biophysics, Chinese Academy of science, Beijing, China). ExpiCHO-S cells were purchased from Thermal Fisher Scientific (Life Technologies Corporation, Carlsbad, CA, USA, code#A29127). Adult healthy Balb/c mice were obtained from Beijing Vital River Laboratory Animal Technology Co., Ltd., Beijing, China. Adult healthy Duhan sheep were obtained from Hengshui Shunyao Sheep Farm (Hebei, China). All experimental procedures using sheep were authorized by the Guide for Animal Care and Use of Laboratory Animals of the Institutional Animal Care and Use Committee of Hebei Agricultural University (Hebei, China; permit number DK596).

### Reagents and kits

PerCP-Cyanine5.5 conjugated CD14 monoclonal antibody (Sa2-8) (code#45014182), PE conjugated CD45 monoclonal antibody (30-F11) (code#12045182), FITC conjugated CD34 monoclonal antibody (RAM34) (code#11034182), APC conjugated CD11b monoclonal antibody (M1/70) (code#17011281), and PerCP-Cyanine5.5 conjugated Ly-6G (Gr-1) monoclonal antibody (RB6-8C5) (code#45593180) were obtained from eBioscience (San Diego, CA, USA). Mouse anti-His monoclonal antibody and HRP conjugated goat anti-mouse IgG were purchased from TransGen Biotech Co., Ltd. (Beijing, China, code#HT501) and Boster Biological Technology Co., Ltd., (Wuhan, Hubei, China, code#BA1050). Non-fat milk was purchased from BD Biosciences (San Diego, CA, USA, code#232100). Restriction enzymes were obtained from New England Biolabs (NEB, Beijing, China).

### RT-PCR and real-time RT-PCR

Total RNA of tissues from heart, liver, spleen, lung, kidney, lymph, uterus, ovary, skin, muscles, breast, and hypothalamus were extracted using the TRIzol RNA isolation system (Life Technologies Corporation, Carlsbad, CA, USA, code#15596018). The first cDNA strand was synthesized using 2 μg of RNA in 20 μl of reaction buffer by reverse transcription using TransScript One-Step gDNA Removal and cDNA Synthesis SuperMix (TransGen Biotech Co., Ltd., Beijing, China, code#AE311-02) following the description of the manufacturer. For the quantitative determination of mRNA levels, real-time RT-PCR was performed on a LightCycler^®^ Nano (Roche, Mannheim, Germany). The PCR reaction mixture in a 20 μl volume containing 10 μl of 2×SYBR Green PCR Master Mix (Roche, Mannheim, Germany, code#06402712001), 2 μl diluted reverse transcriptase product (1:100), and 0.5 μM primer sets. The mRNA levels of the detected genes were calculated after normalization with β-actin. The primers were designed based on the sGCSF sequence (GenBank: XM_001928655.5, XM_015098350.1, and NW_011942481.1) as follows: GCSFv1-F: 5’-GGA CAG ATG CTG TAG CAG G-3’, GCSFv2-F: 5’-CCG GAG GAG CTG GTA CTG-3’, GCSF-R: 5’-GAA GGC CGA CGT GAA GGT-3’, β-ACTIN-PF: 5’-CAG CAG ATG TGG ATC AGC AAG CAG-3’, β-ACTIN-PR: 5’-TTG TCA AGA AAA AGG GTG TAA CGC A-3’. The following RT-PCR thermal cycling conditions were used: after pre-denaturation at 95° for 5 min, 45 cycles were performed at 95° for 10 sec, 53° for 10 sec, and 72° for 15 sec (read), with a melting curve from 60°to 95°at 1°/sec.

### Molecular cloning

The sGCSF full-length DNA samples with stop codons were synthesized by GENEWIZ and amplified with PCR using the following primers that were designed based on sGCSF sequences (GenBank: XM_001928655.5, XM_015098350.1, and NW_011942481.1), in which *Eco*R I and *Nhe* I sites (underlined) were introduced at the 5’ end of forward and reverse primers, respectively. pFUSE-GCSF-wtF: 5’-TCA C<u>GA ATT C</u>GA CCC CCC TTG GCC CTG C-3’, GCSFSP-F: 5’-CTT GTC AC<u>G AAT TC</u>G ATG AAG CTG ATG GGT GAG TG-3’, pFUSE-GCSFH-R: 5’-CCA <u>GCT AGC</u> TTA TCA ATG GTG ATG GTG ATG GTG GGG CTC AGC AAG GT-3’. The following PCR thermal cycling conditions were used: after pre-denaturation at 95° for 5min, 30 cycles were performed at 95° for 15 sec (denaturation), 53° for 15 sec (annealing), and 72° for 20 sec (extension), with a final extension at 72°for 10 min.

### Expression vector construction

The sGCSF DNA that had been PCR amplified was inserted into pRTL1 vector by *Eco*R I and *Nhe* I sites. Positive vectors were identified by colony PCR analysis and were further confirmed by DNA sequencing by GENEWIZ (GENEWIZ, Beijing, China). Furthermore, they were marked as GCSF C-terminal for the mRNA 3’ end domain, as GCSFv1 for the predicted transcript variant 1, and as GCSFv2 for the predicted transcript variant 2.

### Transfection

Transfections were performed with the sGCSF vector following ExpiCHO-S manuals (Life Technologies Corporation, Carlsbad, CA, USA, code#A29127). ExpiCHO-S cells were transfected with full-length pRTL1-GCSF vectors using the ExpiFectamine™ CHO Transfection Kit (Life Technologies Corporation, Carlsbad, CA, USA, code#A29133). Briefly, ExpiCHO-S cells at 1×10^7^ cells/ml (100 ml volume in 500 ml flasks with ExpiCHO-S culture medium) were transfected with 80 μg pRTL1-GCSFV1, pRTL1-GCSFV2, or pRTL1-GCSF C-terminal according to the manufacturer’s protocol. The supernatants were harvested 20 days post-transfection. The recombinant GCSF proteins in the supernatant were detected by dot blot, were purified using High Affinity Ni-Charged Resin (Nanjing Genscript Biotechnology Co., Ltd., Nanjing, China, code#L00223), and detected by SDS-PAGE and Western blot.

### Western blot

Full-length sGCSF proteins in the cell culture supernatant or in the purified soluble sGCSF proteins from culture medium were subjected to 12% SDS-PAGE, were then transferred onto nitrocellulose membranes, and blocked with 5% non-fat milk in PBST (pH 7.4, 0.25% Tween 20) for 2 h. Membrane-associated sGCSF proteins were detected with either mouse anti-GCSF C-terminal antibody or anti-His tag antibody followed by goat anti-mouse IgG-HRP. The results were visualized by staining with the Genshare ECL reagent kit (Genshare Co., Ltd., Xi’an, China, JC-PC001).

### In vitro bioassay

The *in vitro* biological activity of GCSF proteins was determined with a cell proliferation assay with the GCSF dependent cell line M-NFS60. hMCSF (Sino Biological Inc., Beijing, China, code#11792-H08Y) was used as reference standard. Before the assay, M-NFS60 cells were centrifuged at 100 g for 5 min to collect precipitate and re-suspended at a concentration of 1×10^6^ cells/ml in the test medium (RPMI1640 supplemented with 10% fetal bovine serum, antibiotic gentamicin sulfate, and 0.05 mM 2-mercaptoethanol). The test medium (50 μl) was aliquoted into each well of a 96-well tissue culture plate. Both the purified proteins and hMCSF were serially diluted in the test medium. A volume of 50 μl of the diluted protein was added to the wells to achieve concentrations ranging within 0.03-3000 ng/ml. Each protein was tested in triplicate. The cell suspension (50 μl) was added to each well. The plates were incubated at 37° in a 5% CO_2_ atmosphere. After 48 h of incubation, 10 μl of alamarBlue™ HS cell viability reagent (Life Technologies Corporation, Eugene, OR, USA, code#A50101) was added to each well, and the mixture was incubated continually for 3 h under the same conditions. The increased fluorescence was measured (using an excitation between 530-560 nm and an emission at 590 nm) using an EnVision^®^ multimode plate reader (Perkin Elmer, Llantrisant, UK). The biological activity was calculated for each protein from proliferation curves using GraphPad Prism version 6.0.

### In vivo bioassay

Female Balb/c mice (Henan animal research center), 6-8 weeks of age, were used in all animal experiments of this study (de Lichtervelde *et al*. 2012; Zhang *et al*. 2013). The mice were allowed to acclimate for 7 days. Balb/c mice were chosen due to their stimulatory response to GCSF. Animal experiments complied with the Principles of Laboratory Animal Care (Hebei Agricultural University and Zhengzhou Normal University, China) and were approved by the Institute Animal Care and Utilization Committee of the Universities. The treatment groups (n = 3) received a single dose on day 0. For administration via subcutaneous injection (s.c.), 0.5 mg/kg or 10 μg/kg GCSF C-terminal, GCSFV1, GCSFV2, or PBS (as control) were injected. The time of administration was denoted as 0 h. Blood samples were collected at 0, 2, 8, 18, and 30 h from the tail vein, and cells and serums were collected separately. The serums were diluted 100-fold and were analyzed by ELISA using Rabbit polyclonal to 6×His tag^®^ (HRP) (Abcam, USA, code#ab1187) and the QuantaBlu™ NS/K fluorogenic substrate kit (Thermo Fisher Scientific Inc., Rockford, IL, USA, code#15162). The increased fluorescence was measured (using an excitation at 325 nm and an emission at 420 nm) using an EnVision^®^ multimode plate reader. The cells were lysed in an acidic crystal-violet solution (0.1% crystal violet/1% acetic acid, in water). The total white blood cell (WBC) count was manually determined with a NovoCyte 2040R flow cytometer (ACEA Biosciences Inc., San Diego, CA, USA) (Zhang *et al*. 2013). The percentage of polymorphonuclear neutrophils (PMN) among leukocytes was manually determined by using wright stained blood smear glass slides that were examined under an Olympus H7 microscope. The absolute neutrophil count (ANC) was determined by multiplying the total WBC count by the PMN percentage.

### Statistics

The significance of differences between groups was assessed using GraphPad Prism version 6.0 for Windows (GraphPad Software Inc., San Diego, CA, USA) by one-way analysis of variance (ANOVA) with Dunnett’s post-comparison test for multiple groups to control group, or by Student’s t-test (and nonparametric tests) followed by two-tailed comparison tests for two groups.

### Data availability

Strains and plasmids are available upon request. The authors affirm that all data necessary for confirming the conclusions of the article are present within the article and figures.

## Results

### Sequence analysis of sGCSF cDNA

The predicted variant sGCSF sequences and an mRNA 3’ end sequence are available from GenBank. Here, full-length and mRNA 3’ end domains of the sGCSF were compared with those of other animals (Figure 1A and B). The results showed that the sequence of the mRNA 3’ end sequence was consistent with the 3’ end of two prediction variants in GenBank. Comparing the sGCSF cDNA sequences with those of other animals (Figure 1A and Figure S1) showed that the mRNA 3’ end sequence is the same as the GCSFv2 mRNA 3’ end sequence, and the 5’ terminal and the 3’ terminal of GCSFv1 and GCSFv2 are identical with different middle sequences. sGCSF has the highest homology (>91%) with *Bos taurus*, moderate homology (>73%) with human, and low homology with both rattus (>62%) and mouse (>58%). The highly conserved regions were located in the 3’ end domains of GCSFs, suggesting that these 3’ end domains are important for its functions.

**Figure 1.**
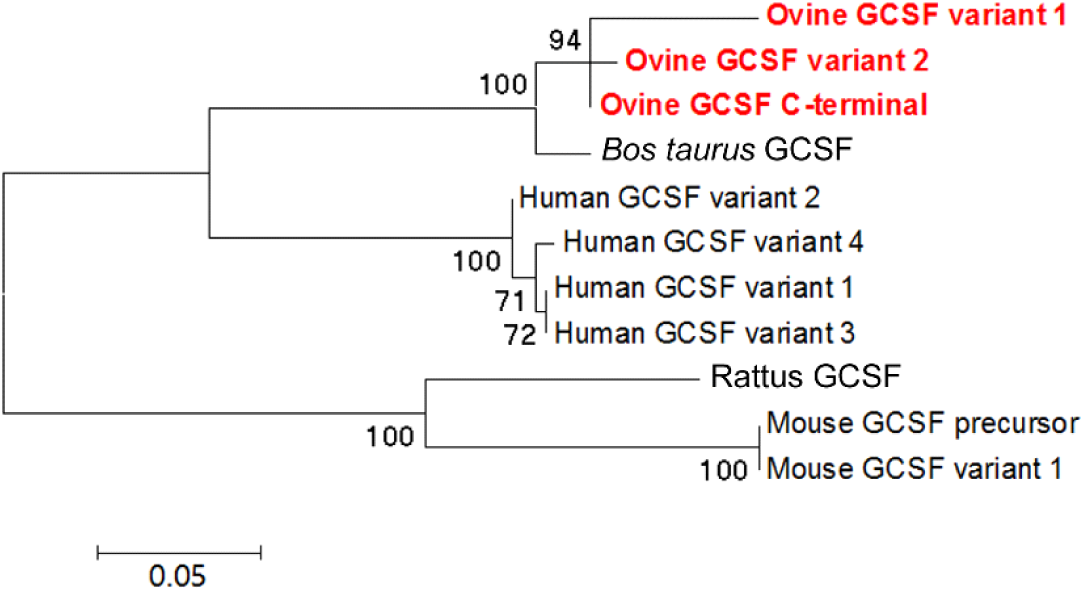
GCSF protein sequences analysis. Molecular phylogenetic analysis of GCSF gene sequences. The evolutionary history was inferred via the Maximum Likelihood method based on a JTT matrix-based model and was assessed in MEGA6 (Tamura *et al*. 2013). The bootstrap consensus tree was inferred from 1000 replicates and was used to represent the evolutionary history of the analyzed taxa.

### GCSF variants expression

Real-time RT-PCR analyses demonstrated that both GCSFv1 and GCSFv2 were expressed at the same time in all tissues at the mRNA level (Figure 2). All amplified lines of GCSFv1 and GCSFv2 were present in the second time of semiquantitative RT-PCR analyses (Figure 2D). As show in Figure 2A and B, Both GCSFv1 and GCSFv2 had significantly lower expression in lung and hypothalamus, and GCSFv1 was significantly highly expressed in breast tissue. These trends were almost stable in other tissues. Quantitative analyses identified higher expression of GCSFv2 than GCSFv1 in all tissues, and the highest expression difference was found in the ovary (Figure 2C).

**Figure 2.**
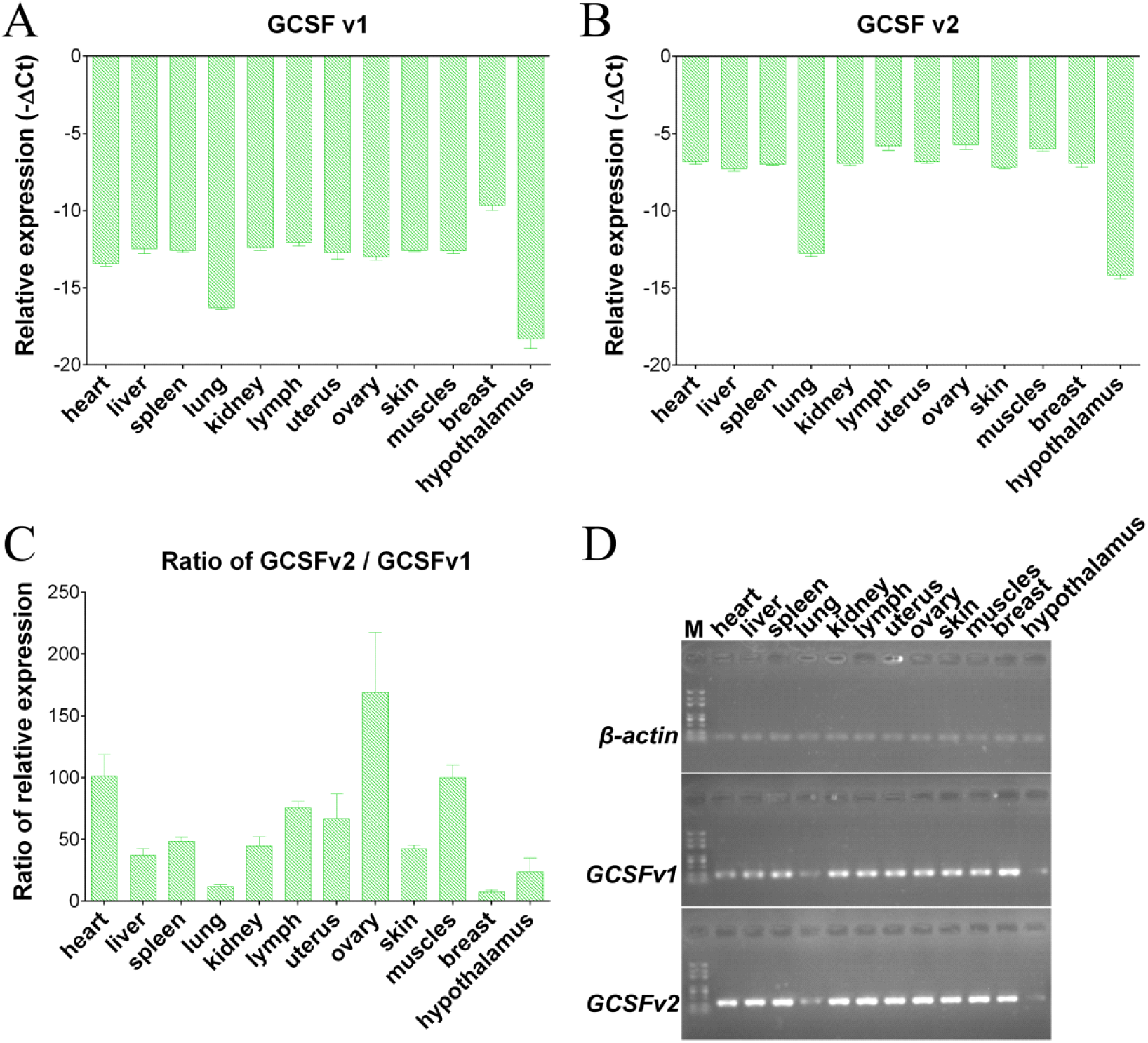
GCSFv1 and GCSFv2 expressions at the mRNA level in different tissues. Quantitative real-time RT-PCR analyses display the relative mRNA levels in different tissues for GCSFv1: β-actin (A), GCSFv2: β-actin (B), and GCSFv2: GCSFv1 (C). (D) Semiquantitative RT-PCR analyses confirm the expression of GCSFv1 and GCSFv2 in different tissues.

### Molecular cloning, sGCSF proteins expression, and purification

The expression plasmids of three types of GCSF proteins (GCSF C-terminal, GCSFv1, and GCSFv2) were expressed in ExpiCHO-S. SDS-PAGE analysis of fractions (Figure 3A) shows one major band of ~90% abundance in each test, with a molecular mass of ~20 kDa, which was visualized by Coomassie blue staining after purification, corresponding to GCSF C-terminal, GCSFv1, and GCSFv2.

**Figure 3.**
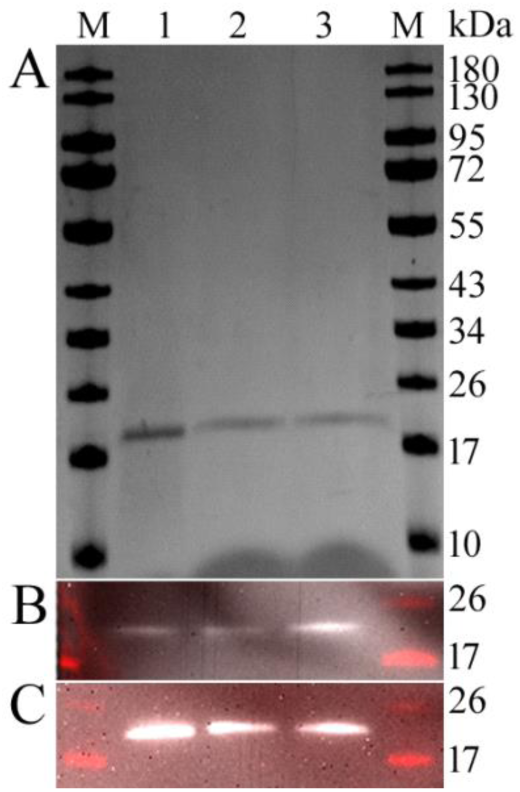
Expression and identification of GCSF proteins. (A) SDS-PAGE of purified recombinant proteins. (B) Recognition of the recombinant proteins by Western blot using anti-6×His tag antibody followed by goat anti-mouse IgG-HRP. (C) Recognition of the recombinant proteins by Western blot using anti-GCSF C-terminal antibody followed by goat anti-mouse IgG-HRP. Lane 1: GCSF C-terminal; lane 2: GCSFV1; lane 3: GCSFV2.

The identities of the secreted proteins were confirmed by using both anti-6×His tag and anti-GCSF C-terminal antibodies via Western blot as illustrated in Figure 3 B and C. Figure 3B shows that the proteins (lanes 1, 2, and 3) were recognized by the anti-6×His tag antibody. Figure 3C shows that the proteins (lanes 1, 2, and 3) were also recognized by an anti-GCSF C-terminal polyclonal antibody.

### In vitro bioassay

The biological activities of the purified proteins were assayed for GCSF activity by determining its ability to stimulate MNFS-60 proliferation. A different amount of protein, which was filtered and normalized for GCSF equivalency, was included in MNFS-60 cell culture medium to replace hMCSF as cell growth factor. As shown in Figure 4, the biological activities of the GCSF C-terminal, GCSFV1, and GCSFV2 were ~44.5/10^th^, 1.03/10^th^, and 6.78/10^th^ of that of the commercial hMCSF, respectively. The EC50 of the hMCSF control was ~2.647 ng/ml, while the EC50 of GCSF C-terminal, GCSFV1, and GCSFV2 were 0.5948, 25.58, and 3.905 ng/ml as a GCSF equivalent. These results showed that GCSF C-terminal activity was best.

**Figure 4.**
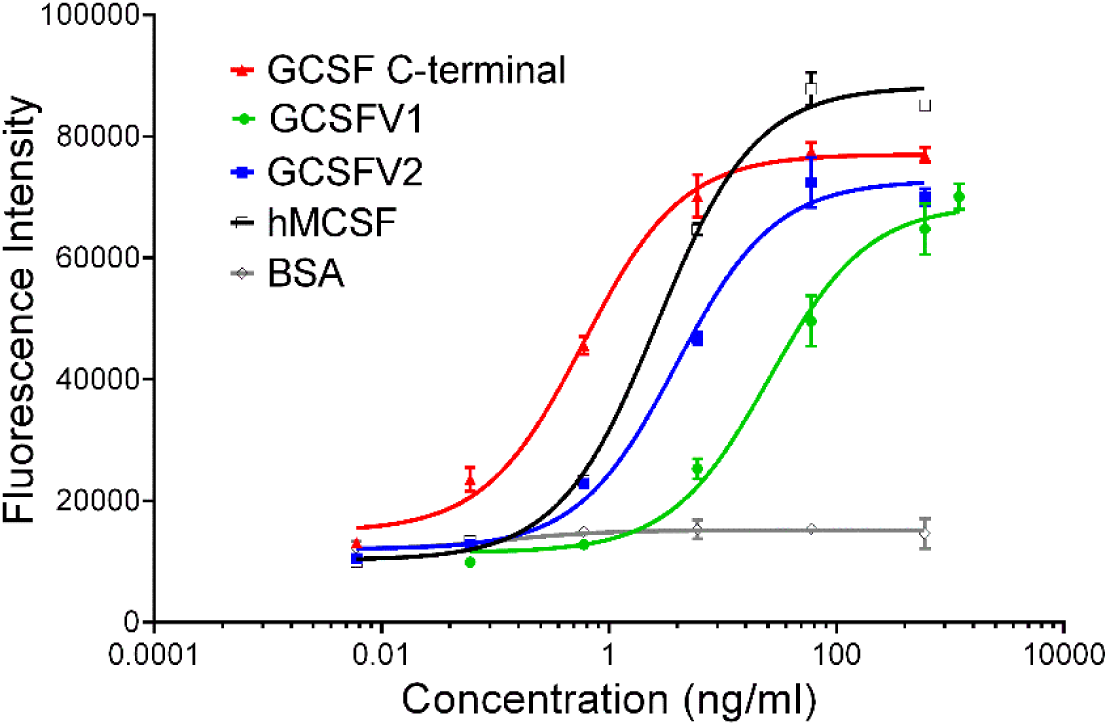
*In vitro* study of the activity of GCSF variant proteins. GCSF proteins stimulate the proliferation of mouse myeloblastic cell line NFS-60 cells in a dose-dependent manner. Cells were treated with various concentrations of GCSF C-terminal (red), GCSFv1 (green), and GCSFv2 (blue). Cell viability was quantified using an AlamarBlue (Invitrogen) assay.

### In vivo studies

Balb/c mice were injected s.c. with 10 μg/kg GCSF C-terminal, GCSFv1, GCSFv2, or PBS control (ctr). The molecular masses of GCSF C-terminal, GCSFv1, and GCSFv2 were 20, 22, and 25 kDa; therefore, the final dosage for each were 50, 46, and 40 nmol/kg. The time-effective curves of proteins were similar. As shown in Figure 5A, all GCSF proteins conferred maximum effect at day 1 (P > 0.1). All GCSFs stimulated increases of CD34+ (progenitors), CD45+ (granulocyte/monocytic lineage), CD11b+ (an integrin characteristic of monocytes and macrophages), and Ly-6G+ (bone marrow cells) cell numbers (Figure 5B) compared to ctr. As shown in Figure 5C, all GCSF proteins exhibited comparable therapeutic effects at the neutrophil level, especially in the GCSF C-terminal group.

**Figure 5.**
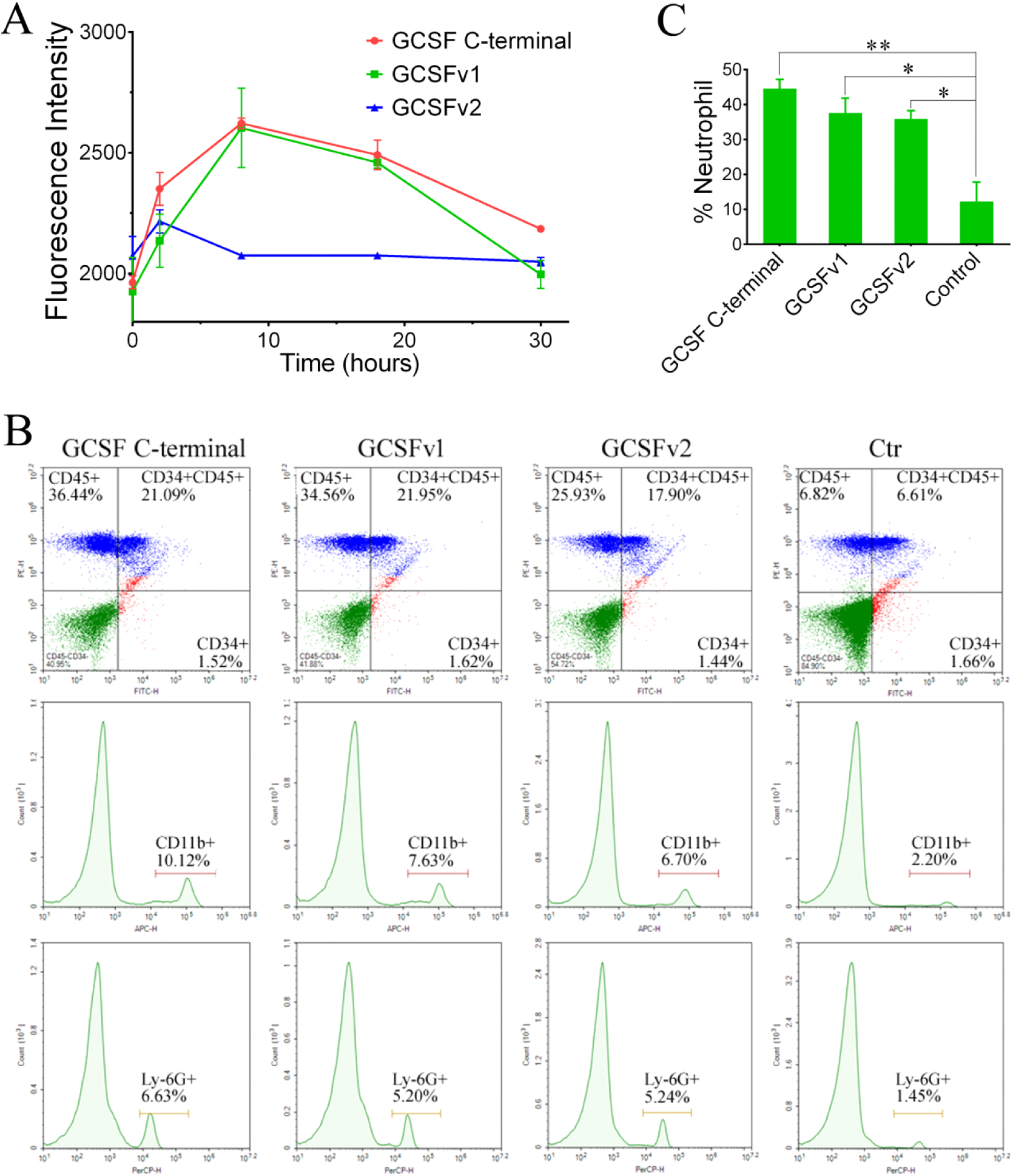
Pharmacokinetics and pharmacodynamics in mice. GCSF C-terminal, GCSFv1, and GCSFv2 were administrated by s.c. injection into Balb/c mice (three per group). (A) Blood was collected between 0 h to 30 h and was analyzed by ELISA using anti-6×His tag antibody. Data were normalized using the maximal concentration during the first 2-8 h. (B) Blood was collected and analyzed for neutrophils by FACS using fluorophore-labeled anti-CD45, anti-CD11b, and anti-Ly-6G antibodies. (C) Representative flow cytometric analyses of mouse neutrophil populations after treatments with GCSF C-terminal, GCSFv1, and GCSFv2. The *p* value of comparison of group Cortrol between group GCSF C-terminal, GCSFv1, or GCSFv2. All statistical analyses were performed using GraphPad Prism 6, and data were averaged across subjects and tested for significance by pair-samples t-test. Bar height denotes the mean average of sample-specific relative affinity values, and values are plotted as means ± SEM from at least three repeats. Asterisks indicate statistical significance as *p* values, \**p*<0.05, \*\**p*<0.001.

## Discussion

In this study, three recombinant fusion proteins, consisting of sGCSF isoforms, were investigated and the results demonstrated that these confer a myelopoietic effect in response to oral delivery in a mouse model. Recombinant human and bovine fusion proteins have been reported to be effective for the development of oral vaccines or drug candidates (Eapen *et al*. 2019; Nguyen 1994; Stephens *et al*. 2019; Takehara *et al*. 2019; Wu *et al*. 2019). However, this report is the first demonstration that sheep GCSF isoform proteins assume the same roles.

Human and bovine GCSFs have gained clinical use due to their ability to selectively stimulate proliferation and differentiation of progenitor cells and to activate the maturation of neutrophils (Caselli *et al*. 2016). There has been no report on sheep GCSF function except for the predicted gene sequences in GenBank. Here, the two predicted GCSF variants GCSFv1 and GCSFv2 were cloned, and their constant segments were marked as GCSF C-terminal. All three proteins had the same function as human and bovine GCSF in this experiment. The activity and mRNA expression level of GCSFv2 was higher than that of GCSFv1. Their constant part at the GCSF C-terminal has even more activity than both. Pharmacokinetics with GCSF C-terminal, GCSFv1, and GCSFv2 were administrated by s.c. injection into Balb/c mice, which showed that the maximal concentration could be detected within the first 2 h in the GCSFv2 group, and within the first 8 h in GCSF C-terminal and GCSFv1. This suggests that GCSFv2 is the short-term agent, while GCSFv1 is the long-term agent, due to its shorter circulation half-life and more activity compared with GCSFv1. Furthermore, their short circulation half-life problem also had to be modified as human and bovine GCSF. The Fc fusion format was applied to generate a GCSF dimer, which exhibited biological activity *in vivo* (Yan 2016). The same method can be used for sGCSF.

The ability to stimulate cell proliferation *in vitro* was higher for GCSF C-terminal molecules compared with both GCSFv1 and GCSFv2. The *in vitro* biological activity of the GCSF C-terminal was about 7-50-fold higher than the activities of both GCSFv1 and GCSFv2, which suggests that C-terminal plays an important role in the protein function. Moreover, the N-terminal has a spatial interference impact on the GCSF interaction with the receptor. Biological activity assays in healthy mice demonstrated that the GCSF C-terminal molecule offers advantages over GCSFv1 and GCSFv2. The GCSF C-terminal showed a circulation half-life of 8 h, which was identical to those of GCSFv1 and GCSFv2. The *in vivo* response, which comprised of the ability of GCSF to stimulate neutrophil release, was more pronounced for GCSF C-terminal compared with GCSFv1 and GCSFv2. After 24 h of a single subcutaneous injection of the GCSF C-terminal, mice exhibited a 3-5-fold increase of circulating neutrophils compared with ctr group, albeit with a larger margin of error. These results indicated that the GCSF C-terminal could be used as a candidate for drug development and as a target of molecular genetic breeding. However, the half-life of the proteins and the sustained duration effect need to be increased and this approach needs to be further developed for other protein drugs and therapeutic applications.

## Conclusion

The two ovine predicted GCSF variants GCSFv1 and GCSFv2 were cloned. The activity and expressed mRNA level of GCSFv2 were higher than those of GCSFv1 and it showed the highest homology with *Bos taurus*. The protein function of the GCSF C-terminal was identical to that of both GCSFV1 and GCSFV2 proteins, which are stably expressed in mammalian cells. The activity of the GCSF C-terminal was best. These findings provide an approach for the future development of orally efficacious protein drugs, and provide a candidate target for the future development of sheep genetic breeding.

## Acknowledgement

This work was supported by funding from the National Key RD Program of China [2018YFD0502100]; National Modern Agricultural Industry Technology System Construction Project of China [CARS-38] and [CARS-39].

We thank Feng Wang from Institute of Biophysics, Chinese Academy of science in the help of material providing and experiment advices. We thank Wenxin Cao, Sisi Ju, xuejiao Yin, Kun Gao, and Jiwei Zhang from College of animal science and technology, Hebei Agricultural University in the help of operation on test with sheep.

